# Nitrate reductase activity is required in *Medicago truncatula*-*Sinorhizobium meliloti* nitrogen-fixing symbiosis

**DOI:** 10.1101/2024.05.31.596865

**Authors:** Bosseno Marc, Demba Alexandre, Horta Araújo Natasha, Colinet Dominique, Pacoud Marie, El Fazaa Yassine, Lepetit Marc, Clément Gilles, Brouquisse Renaud, Boscari Alexandre

## Abstract

Nitrate reductase (NR) is a key enzyme in higher land plants, catalyzing the rate-limiting reduction of nitrate to nitrite in the nitrate assimilation pathway. Phylogenetic analysis of NR protein sequences indicates that duplication events responsible for the existence of two NR branches, corresponding to NR1 and NR2 genes, occurred after the divergence of the different orders within the Rosids clade. A third NR sequence branch, named NR3-type, emerged in the inverted repeat-lacking clade of the Fabales order. An intriguing feature of the NR3-type sequences is the absence of conserved phosphorylation sites in the two hinge regions, in contrast to all other NRs. To investigate the respective roles of *MtNR1*, *MtNR2* and *MtNR3* in *M. truncatula*, three single *Tnt1* retrotransposon-tagged *nr* mutants and one *nr1/nr2* double mutant were analyzed on plants growing either on nitrate, or during the nodulation process. Overall, the absence of phenotypes observed in *M. truncatula* single mutants suggests a significant functional redundancy between the different NRs in *M. truncatula*. The most striking outcome of this work is the almost complete impairment of nodulation capacity observed in the *nr1/nr2* double mutant, demonstrating that NR activity is required for the functioning of the N_2_-fixing symbiosis.

## Introduction

In plants, yeasts, algae, and fungi, nitrate reductase (NR) is a key enzyme for nitrogen acquisition. The enzyme catalyses the reduction of nitrate (NO_3_^-^) to nitrite (NO_2_^-^). Nitrite is then reduced to ammonia (NH_4_^+^) by nitrite reductase (NiR) before being assimilated into the amino acids and nitrogen compounds of the cell (Campbell, 1988; Meyer and Stitt, 2001). NR is a homodimer consisting of two approximately 100 kDa polypeptide chains. It is responsible for the first intracellular and rate-limiting step in nitrate assimilation. Each chain binds three cofactors in individually folded domains and the enzyme operates as an internal electron transport chain (Campbell, 1988). The C-terminal domain contains a flavin adenine dinucleotide (FAD) cofactor and accepts two electrons from NADH or NADPH. It then transfers these electrons to the middle domain, which carries a b5-type cytochrome heme. The electrons are then transported to the catalytic site, which contains a molybdenum cofactor (Moco) in the N-terminal domain, where substrate reduction takes place. In addition to the regulation of steady-state levels of NR protein at the transcriptional and protein turnover levels (Hoff et al., 1994), post-translational modification of NR by protein phosphorylation is an important regulatory mechanism (Douglas et al., 1995; Bachmann et al., 1996; Su et al., 1996; Lillo et al., 1997; Lillo et al., 2003).

In the Rosid clade, leguminous plants have developed a mutually beneficial relationship with soil bacteria (rhizobia) that enables them to convert atmospheric nitrogen (N_2_) to ammonia (NH_4_^+^) through the activity of nitrogenase (Postgate, 1982). In exchange for NH_4_^+^, plants provide an ecological niche to the bacteria for their development and carbon nutrients for their functioning in newly formed organs called nodules (Udvardi and Day, 1997; Terpolilli et al., 2012). In legume nodules, the bacterial nitrogenase produces NH_4_^+^ instead of the NR-NiR pathway and supplies it to the plant. However, numerous studies have shown high levels of NR expression and activity in symbiotic nodules (Streeter, 1985a; Streeter, 1985b; Arrese-Igor et al., 1990; Silveira et al., 2001), and the localization of NR mRNA within the infected regions of pea root (Kato et al., 2003) and *M. truncatula* (Horchani et al., 2011) nodules raises the question of what NR can do in the N_2_-fixing symbiosis, since reduced nitrogen is provided by nitrogenase. Despite the identification of NR mutants in legumes (Feenstra and Jacobsen, 1980; Ryan et al., 1983), no study fully elucidated the significance of these enzymes in the process of nitrogen-fixing symbiosis. Previous works showed that NR activity is required for energy metabolism and N_2_ fixation in *M. truncatula* mature nodule (Horchani et al., 2011; Aridhi et al., 2020; Chammakhi et al., 2022). Recently, Berger et al. (Berger et al., 2020a) showed that, during the symbiosis between *M. truncatula* and *Sinorhizobium meliloti*, *MtNR1* and *MtNR2* expression peaks coincide with that of NO production, and that these NO production peaks are abolished by either the use of tungstate, a NR inhibitor, or by a NR1/NR2 RNAi knock-down strategy. These results point out the involvement of NR in the regulation of NO during the symbiotic process. The presence of three NR genes in the *M. truncatula* genome (Puppo et al., 2013) raises the question of their respective roles in nitrate-fed as well as in symbiotic *M. truncatula* plants.

In the present work, a phylogenetic analysis was first performed to understand the evolutionary relationships among NR protein sequences belonging to different orders of the Rosids clade. Then, to investigate the respective role of *MtNR1*, *MtNR2* and *MtNR3* in *M. truncatula*, we selected *Tnt1* retrotransposon-tagged *Medicago nr* mutants, and generated a double nr1/nr2 mutant. These different lines were subsequently analysed on plants growing on nitrate and during the nodulation process.

## Materials and methods

### Plants growth and inoculation conditions

Seeds of *M. truncatula* cv R108 were scarified, surface-sterilized and germinated as in del Giudice et al. (2011). Germinated seedlings were transferred either to new plates with Fahräeus medium and 0.2 mM NH_4_NO_3_, to individual pots, or to hydroponic systems. Plants in individual pots (Pots PP polypropylene, 9*9*9cm, Soparco), containing a mixture of vermiculite and perlite (2:1, v/v), were watered every 2-3 days, two times with water for one time with 1 g.l^−1^ of nutritive solution. For hydroponic culture, 6 day-old plants were transferred into tanks containing an aerated basal nutrient solution HY (Lambert et al., 2020) supplemented with 1 mM NH_4_Cl as an N source assimilable in nitrate reductase-deficient plants to sustain plant growth for one week. Nutrient solution was then renewed and either supplemented with 1 mM of KNO_3_ or inoculated with *S. meliloti* (10^7^ cfu ml^−1^) plus 10 µM of KNO_3_. Nutrient solution was renewed every week. Plants were grown in climatic chamber (20-23°C with a 16h light/8h dark photoperiod). Plants in plates and pots were inoculated five days after transfer with *S. meliloti* 2011 strain at OD600 0.5 as described in del Giudice (2011).

*A. M. truncatula* homozygous *Tnt-1* insertion mutants in R108 genetic background, NF15469 (*Tnt1* insertion in *MtNR1)*, NF14768 (*Tnt1* insertion in *MtNR2*) and NF11190 (*Tnt1* insertion in *MtNR3*) were obtained from the Noble Research Institute (Ardmore, OK, USA) (Tadege et al., 2008). The oligonucleotide primer sets (Table S1) were designed to allow screening of the intact *MtNR* genes and were also used in combination with LTR4 and LTR5 oligonucleotide primers (Ratet et al., 2010) to confirm *Tnt1* boundaries within three *MtNR* genes. The NF15469 (with 6 *Tnt1* insertions), NF14768 (with 5 *Tnt1* insertions), and NF11190 (with 20 *Tnt1* insertions) were backcrossed 3, 3, and 4 times, respectively, to clean up the genetic background of these mutants following the protocol described by Bosseno et al. (2020). *Mtnr1* and *Mtnr2* lines have been crossed to create a double mutant. Despite numerous attempts with different nitrogen sources (ammonia, urea, etc.), it has not been possible to get to the point of flowering and seed production stage in the double mutant seedlings knock-out (KO) on *NR1* and *NR2* gene. The characterization of the *Mtnr1/nr2* mutant was carried out using segregated seeds from crosses of the two single mutants, KO for *MtNR1* and heterozygous for *MtNR2*.

### Phylogenetic analysis of NR protein sequences

Nitrate reductase (NR) protein sequences were initially obtained from BLASTP searches against the refseq_protein database at NCBI (www.ncbi.nlm.nih.gov/) using *M. truncatula* NR1, NR2, and NR3 sequences as queries and an e-value threshold of 1.0e-5. BLASTP searches were restricted to Rosids (taxid:71275), excluding *Medicago* (taxid:3877), with a maximum number of hits of 500 on the one hand, and to Poales (taxid:71275) with a maximum number of hits of 100 on the other. The resulting sequence dataset was supplemented with NR protein sequences from Rosids obtained from an exhaustive manual search of the Phytozome genomics database (www.phytozome.jgi.doe.gov), and of the UniprotKB (www.uniprot.org) and NCBI non-redundant (www.blast.ncbi.nlm.nih.gov) protein databases. A CD-HIT analysis was then performed with a similarity threshold of 95% to remove redundancy (Li and Godzik, 2006). Finally, orders belonging to Rosids that contained only one sequence was eliminated and only one species were retained for each genus. A total of 121 non-redundant protein sequences, ranging from 582 to 956 amino acids and belonging to 64 species were retained for phylogenetic analysis and are listed in Table S2.

The 121 NR protein sequences were aligned using MAFFT (v7) with the --auto option, which automatically selects the appropriate alignment strategy based on data size (Katoh and Standley, 2013). Poorly aligned regions were removed using trimal (v1.4) with the - automated1 option, which is optimized to automatically select the best trimming method for maximum likelihood (ML) phylogenetic analysis (Capella-Gutiérrez et al., 2009). The ML phylogeny of NR protein sequences was inferred using IQ-TREE (v2) (Minh et al., 2020) with automated model selection (Kalyaanamoorthy et al., 2017). Support values were based on a Shimodaira–Hasegawa approximate likelihood ratio test (SH-aLRT) (Guindon et al., 2010) combined with an ultrafast bootstrap approximation (UFboot) with 1,000 replicates each (Hoang et al., 2018). Only support values greater than or equal to 80% and 95% for SH-aLRT and UFboot, respectively, were considered. Phylogenies were visualized and annotated using iTol (Letunic and Bork, 2021).

To investigate the absence of NR1 in some species of the IRLC clade, a protein-to-genome comparison using Exonerate (v2) with a score threshold of 500 (Slater and Birney, 2005) was performed with the three *M. truncatula* NR protein sequences in the genomes of *Pisum sativum* (NCBI accession number GCF_024323335.1), *Vicia faba* (NCBI accession number GCA_948472305.1) and *Cicer arietinum* (NCBI accession number GCF_000331145.1).

### Analysis of Fabales sequence phosphorylation sites

The 46 NR protein sequences from *Fabales* and *Cucurbitales* species, as well as *A. thaliana,* were aligned using MAFFT (v7) with the --auto option, which automatically selects the appropriate alignment strategy based on data size (Katoh and Standley, 2013). The amino acids in the hinge regions between Moco and Heme and Heme and FAD were identified using the PFAM tool (El-Gebali et al., 2019). Prediction of the phosphorylation sites within each sequence included in the alignment was conducted using MusiteDeep ((Wang et al., 2017); https://www.musite.net). Phosphorylation sites with a probability higher than 0.5 were selected and compared with predictions generated by the PlantPhos tool (Lee et al., 2011). Predictions and *A. thaliana* known phosphorylation sites were manually annotated in the alignment using Microsoft Word.

### Gene expression analysis

Harvested nodules and roots were frozen in liquid nitrogen and ground. Total RNA extraction was performed using the RNAzol ® RT reagent following the manufacturer’s recommendations (Sigma-Aldrich, USA). After the DNAse treatment with RQ1 DNAse (Promega, USA), 1 µg or total RNA was used for the cDNA synthesis (GoScript ™ Reverse Transcription System, Promega). Quantitative real-time PCRs were performed in the Agilent AriaMx System using the GoTaq® qPCR Master Mix (Promega, USA). Two reference genes (MtC27 and a38) were used to normalize the transcript level. The complete list of primers is reported in Table S1. Data were quantified using the Aria Software and analyzed with RqPCRBase, an R package working on R computing environment for analysis of quantitative real-time PCR data (Rancurel et al., 2019).

### Nitrogen-fixing capacity

The nitrogen-fixing capacity of nodules was determined *in vivo* by measuring the reduction of acetylene to ethylene, as previously described as acetylene-reducing activity (ARA, (Hardy et al., 1968)). Nodulated roots were harvested and incubated at 30 °C for 1 h in rubber-capped tubes containing a 10% (v/v) acetylene atmosphere. Two biological replicates have been performed with five technical replicates. Ethylene concentrations were determined by gas chromatography (Agilent GC 6890N, Agilent Technologies, Santa Clara, CA, USA) equipped with a GS-Alumina (30 m × 0.534 mm) separating capillary column.

### Metabolomic

Mature nodules (4 wpi) were collected in three replicates from *M. truncatula* cultures in pots. Metabolites were extracted and analyzed by GC/MS after grinding and homogenization in liquid nitrogen as previously described (Clément et al., 2018; Lambert et al., 2020).

### Measurement of NO production

NO detection was performed as in Horchani et al. (2011) using the 4,5-diaminofluorescein probe (DAF-2, Sigma-Aldrich) with the following changes. Either nodules (20-30 mg fresh weight) or root segments (50-100 mg fresh weight) were incubated in 1 ml of detection buffer (10 mM Tris-HCl pH 7.4, 10 mM KCl) in the presence of 10 µM DAF-2. The production of NO was measured with a spectrofluorimeter-luminometer (Xenius, SAFAS, Monaco).

### Measurement of Nitrate Reductase Activity

Tissue samples are ground with mortar and pestle in liquid nitrogen. The total proteins are extracted from 100 mg of powder using the following extraction buffer: 25 mM Tris HCl pH 8.5, 1 mM EDTA, 20 μM FAD, 0.04% (v/v) Triton, 10 μM NaMO4, 1 mM DTT, E64 μM, 2 mM PMSF. The extracts are centrifuged (15000 g, 15 min). Nitrate reductase activity was assayed at 28°C by measuring NO_2_^-^ production as described in Horchani et al. (2011).

### Statistical Analysis

The non-parametric Mann–Whitney test was used for two groups of variables. For three or more than three groups of variables, the Kruskal–Wallis and Dunn test were applied using the Past3 software (https://folk.uio.no/ohammer/past/).

## Results

### Phylogenetic analysis of NRs proteins sequences belonging to Rosids clade

Maximum likelihood phylogenetic analysis was performed to understand the evolutionary relationships among NR protein sequences belonging to 10 different orders of the Rosids clade (Fig. 1 and Fig. S2). NR sequences from the Poales clade were used as an outgroup to root the phylogenetic tree. The phylogeny was constructed from 121 different NR protein sequences from 64 species (Table S2), including the well-characterized NIA1 and NIA2 from *A. thaliana* and NR1, NR2, and NR3 from *M. truncatula* (Fig. 1 and Fig. S2). In the Rosids clade, at least one species with two or more NR protein sequences were identified for each order except for the Sapindales. With the exception of the Myrtales, each order formed a monophyletic group, suggesting that the duplication events leading to the emergence of distinct NR genes occurred subsequent to the divergence of the different orders within the Rosids clade as indicated by the red dot on Fig. 1 and Fig. S2.. In the Fabales and Brassicales, where NR protein sequences are well characterized for *M. truncatula* and *A. thaliana*, respectively, an ancestral duplication event is observed at the origin of NR1 and NR2 in *M. truncatula* and NIA1 and NIA2 in *A. thaliana* (Fig. 1). Within the Fabales order, this ancestral duplication event occurred after the separation with the *Mimosoids* clade (Fig. 1). Consequently, this clade is phylogenetically more distanced from the other clades included in the analysis (Wojciechowski et al., 2004). In all clades of the Fabales except for Robinoids clade’s *L. japonicus*, both NR1 and NR2 orthologs have been identified. Instead *Lotus japonicus* possesses a single NR sequence closely related to MtNR2 (Fig. 1). Furthermore, a more recent duplication event of the NR1 gene occurred in the ancestor of the Inverted Repeat-Lacking Clade (IRLC) of the Fabales (Fig. 1). This particular duplication event gave origin to the NR3 gene found in all species of the IRLC clade studied to date. Interestingly, however, the presence of both NR1 and NR3 sequences was found only in *M. truncatula* and the related *Trifolium subterraneum* (Fig. 1). Only NR3 sequence was found in the other species within the IRLC clade, namely *Pisum sativum*, *Vicia faba* and *Cicer arietinum*, suggesting a potential loss of the NR1 gene in these species. Accordingly, a search for orthologs of the three *M. truncatula* genes in the genomes of *P. sativum*, *V. faba* and *C. arietinum* yielded results only for NR2 and NR3. Interestingly, two ancestral duplication events occurred in the *Cucurbitales,* giving rise to three NR sequences, suggesting that orders beyond the Fabales may harbor additional types of NR. Such phenomena are not exclusive to species capable of nitrogen-fixing symbiosis. Recent duplications have also been noted across various orders, particularly in species with large genomes like *Glycine max,* resulting in an increased number of closely related NR paralogs (Fig. 1).

**Figure 1:**
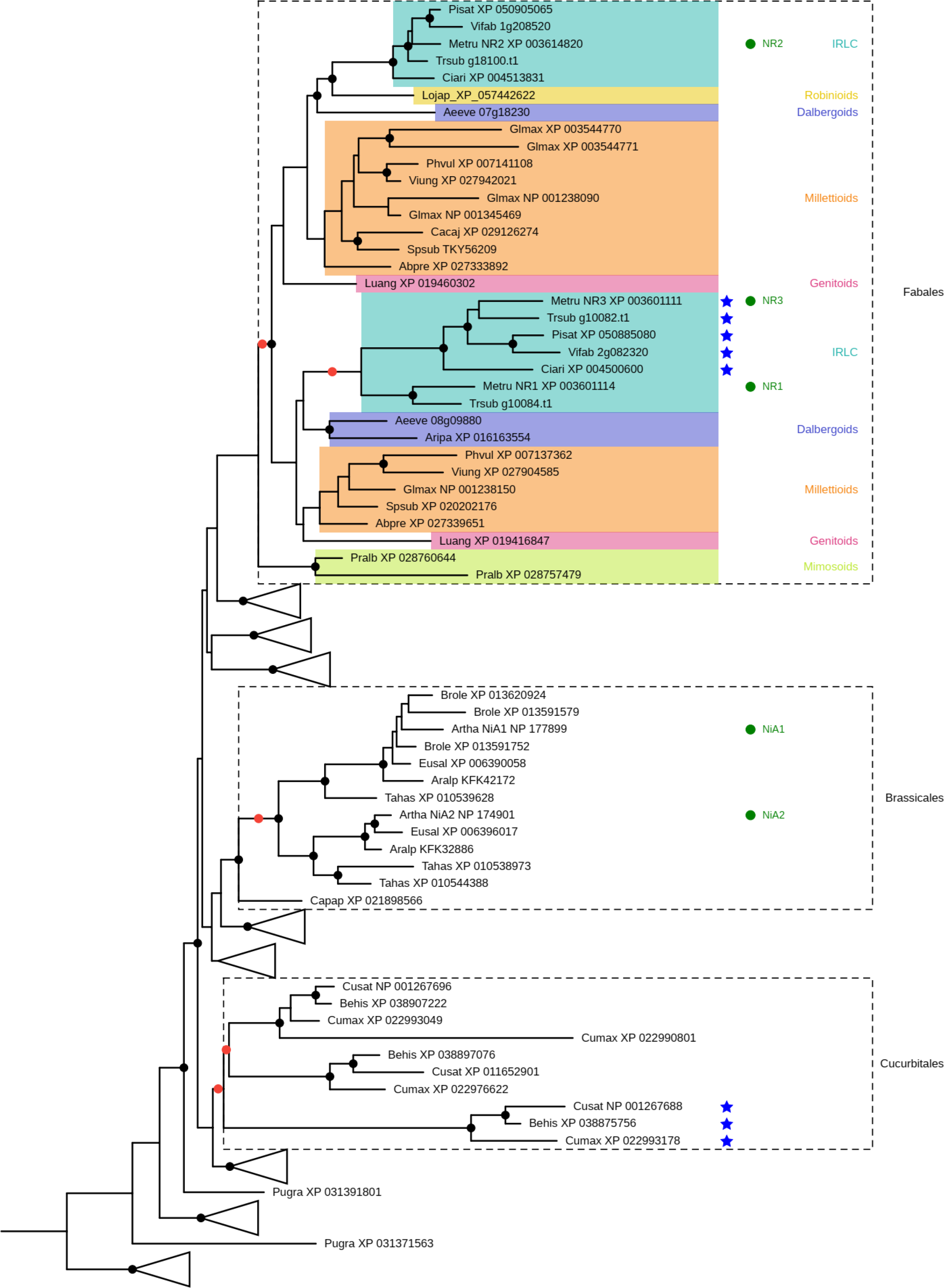
**Simplified representation of the maximum likelihood phylogenetic tree of 121 NR protein sequences from the Rosids clade and the Poales clade, which was used as an outgroup to root the phylogenetic tree**. The well-characterized NIA1 and NIA2 from *A. thaliana* and NR1, NR2, and NR3 from *M. truncatula* are indicated by a green dot. NR sequences with potential loss of phosphorylation site are indicated by a blue star. Red dots on selected nodes indicate the duplication events at the origin of the two or three groups of NR protein sequences found in the Brassicales, Fabales, and Cucurbitales.

### Analysis of the phosphorylation site in NR hinge regions of *Fabales*

Nitrate reductases are molybdoenzymes that function as dimers or homotetramers, where each subunit of this protein complex contains 3 prosthetic groups: the cofactors flavin adenine dinucleotide (FAD), the molybdenum cofactor (Moco), and the heme (Campbell, 2001) (Fig. 2A). The three NR proteins of *M. truncatula* exhibit a significant level of similarity (>70%) (Berger et al. 2020. Fips). However, on a multiple protein sequence alignment involving representatives from the *Fabales* and *Cucurbitales* orders, as well as the two well-characterized *A. thaliana* NR sequence, notable differences emerged within two hinge regions (Fig. 2B and C). The prosthetic groups are bound to distinct protein domains interconnected by flexible hinge regions notably known for their role in post-translational regulation in response to different signals such as light intensity, CO_2_ or O_2_ levels (Kanamaru et al., 1999; Kaiser and Huber, 2001). In *A. thaliana*, Serine 534 in Hinge 1, situated between the Moco and heme domains and Serine 627 in hinge 2, located between the heme and FAD domains, are associated with specific post-translational regulation mechanisms through phosphorylation (Lea et al., 2004; Wang et al., 2010).

**Figure 2:**
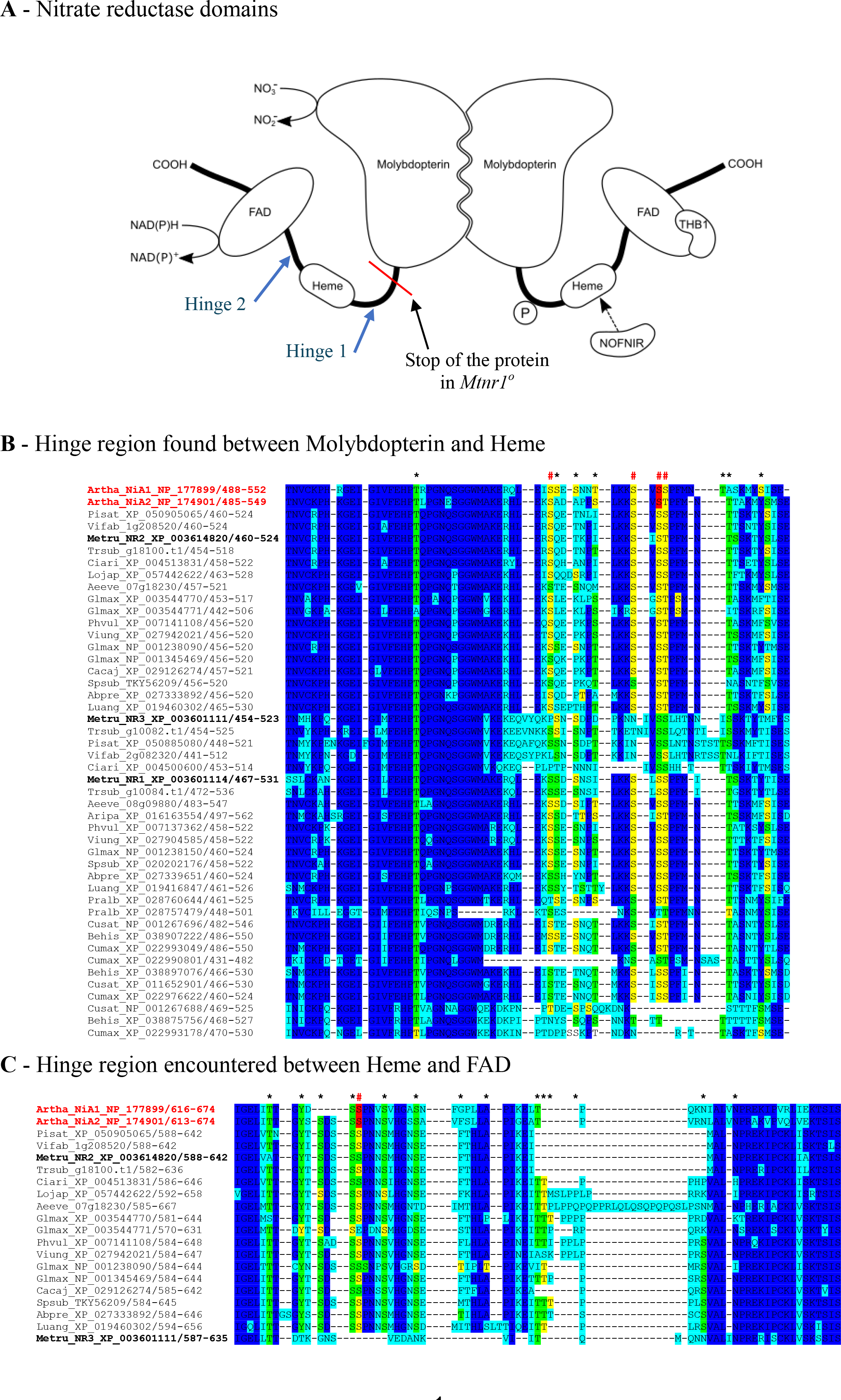
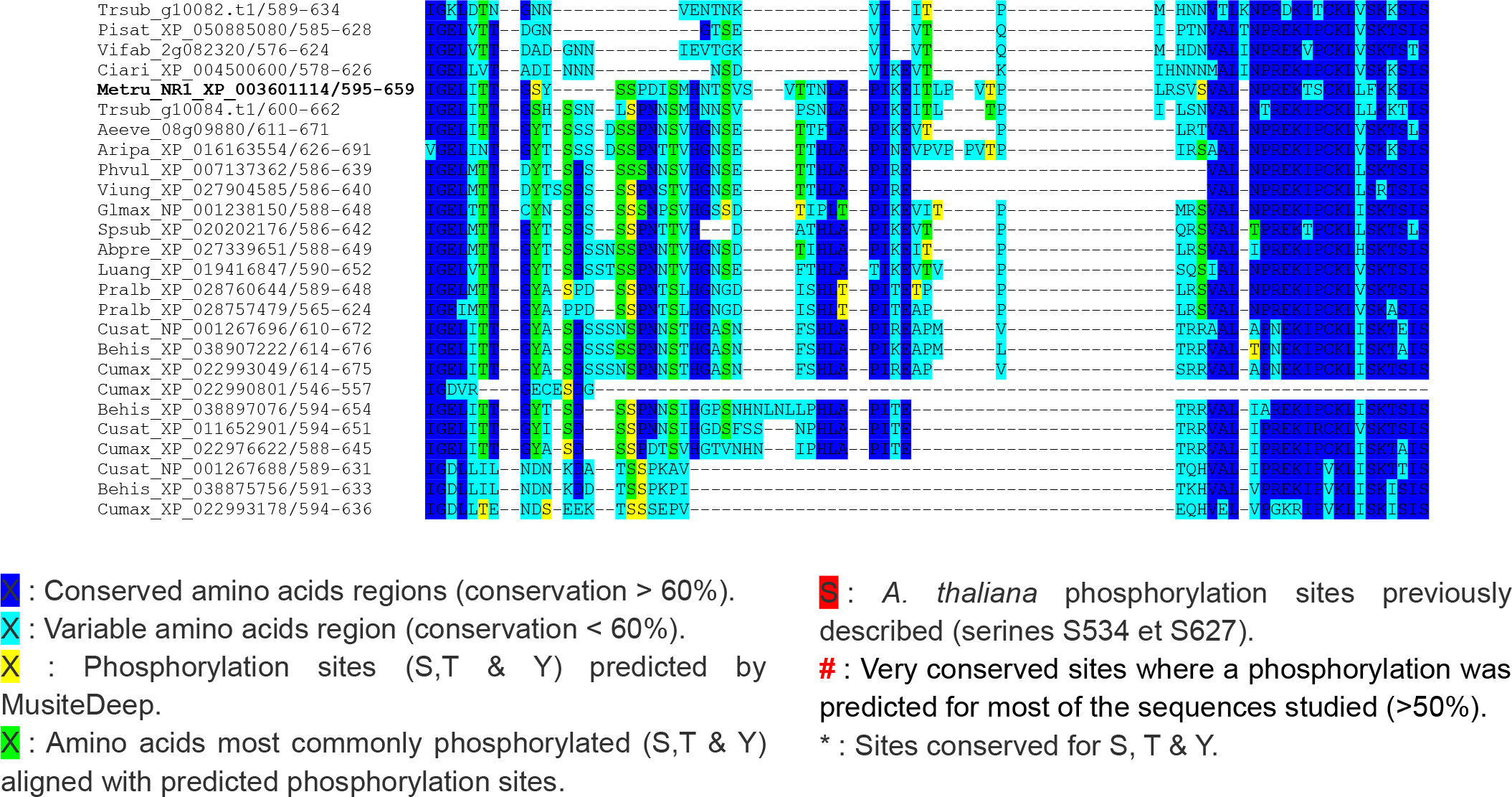
**The two NR hinge regions and their potential phosphorylation sites.** (A) Schematic representation of plant Nitrate Reductase (NR) structure. Each subunit of this protein complex contains 3 prosthetic groups: the cofactors flavin adenine dinucleotide (FAD), the molybdenum cofactor (Moco), and the heme. NR has two "hinge" regions situated between the Moco and heme domains (Hinge 1) and between heme and FAD domains (Hinge 2). Phosphorylation site in hinge 1 is represented. (B) Alignment of the NRs protein sequences of *A. thaliana*, *Cucumis sativus* and NRs belonging to the Fabales family in region between the Moco and heme domains, (C) and between the heme and FAD domains. Sequences are labeled with codes explained in Table S2. *A. thaliana* is shown in black, while the names of *Fabales* sequences appear in green or orange, depending on the type of nodule formed by the plant: determinate or indeterminate, respectively.

The multiple protein sequence alignment revealed that Serine 534 in hinge 1 is conserved among all protein sequences from Fabales used for the alignment (Fig 2B), with the exception of five NR3 sequences (MtNR3, Ciari_XP004500600, Pisat XP050885080, Vifab 2g082320, and Trsub g10082.t1). Similarly, the second phosphorylation site, serine 627, in the hinge 2 region, also exhibited non conservation of the serine amino acid among these five NRs, in contrast to the other sequences (Fig. 2C). Additionally, the analysis of NR sequences using a tool that predicts global phosphorylation sites, such as the Musite tool (https://www.musite.net/), confirmed the lack of predicted phosphorylation sites for the aforementioned sequences (Fig. 2B and C). The lack of conserved of post-translational regulatory sites in all proteins phylogenetically close to MtNR3 suggests a distinct mechanism of regulation for NR3-type NR, potentially important within the IRLC clade. Interestingly, a lack of predicted phosphorylation sites was also evident for a group of three NR sequences in *Cucurbitales* (Cusat NP001267688, Behis XP038875756, and Cumax XP022993178), suggesting a convergent loss of this regulatory motif in specific NR sequences of Fabales and *Cucurbitales* through independent evolutionary events in two distantly related orders within the *Rosids* clade (Fig. 2B and C).

Given that *MtNR3* from *M. truncatula* is exclusively expressed during symbiotic interaction and particularly at the meristem level of the nodule primordium (Berger et al., 2020a), we sought to determine if other *NR3-*type *NRs* from IRLC clade exhibited similar expression pattern to *MtNR3*. We conducted a database search to collect information regarding the expression profile of the three *NR3*-type genes (Fig S2). In *Cicer arietanum*, the expression of *CaNR3* is notably exceeded that of *CaNR2* in most of the analyzed plant organs, as depicted in Figure S2A. Interestingly, *CaNR2* expression was exclusively detected in roots and nodules, as shown in Figure S2A. Meanwhile, in *Pisum sativum*, *PsNR2* expression predominantly occured in leaves and roots, while *PsNR3* was mainly expressed in pods, seeds, and nodules (Fig S2B). Regrettably, no information was found regarding the expression profile of the two NRs in *Vicia faba*. In contrast to *MtNR3*, *PsNR3* and *CaNR3*, two additional NR3-type NR, are not exclusively expressed during nodulation. Consequently, the hypothesis that these NR3s have a specific regulation in relation to nodule function has been ruled out.

#### Function of *M. truncatula* NRs in plants grown on nitrate source

The function of nitrate reductase is to reduce nitrate into nitrite. Nitrite will be further reduced to NH_4_^+^ by NiR to fuel the amino acid metabolism through the action of GS/GOGAT. In addition to this nutritional function, NO_2_^-^ is fuels the NO synthesis (Crawford and Campbell, 1990; Pelsy and Caboche, 1992). The presence of three genes in the *M. truncatula* genome encoding distinct NR isoforms may suggest that they have distinct roles either in N nutrition or in NO production. To investigate the respective role of *MtNR1*, *MtNR2* and *MtNR3* in *M. truncatula*, we selected *Tnt1* retrotransposon-tagged *Medicago* mutants. We identified a *Tnt1*-insertion lines for each *NR* gene of *M. truncatula* (Fig S2). The three *NR* genes are composed with four exons (Fig. S3). In NF14768 and NF11190 lines, *Tnt1* is inserted in the first exon of *MrNR2* and *MtNR3* genes respectively, likely resulting in the inactivation of the gene and the arrest of synthesis of the respective proteins. In NF15469 line, *Tnt1* is inserted in the fourth exon of *MtNR1*. In theory the production by the mutant of a shortened NR1 protein of 480 amino acids (compared to the 902 amino acids of the wild-type protein) cannot be excluded. However, because this hypothetical protein would retain only the Molybdopterin domain and would lack both the heme and the FAD domains, known to be essential to the NR catalytic function, this hypothetical protein is also most probably not functional (Fig. S3, Fig. 2A). To validate further these lines, the total NR activity of *Mtnr1*, *Mtnr2* and the segregating double mutant *Mtnr1/nr2*, were analysed using hydroponic culture supplemented with 1 mM KNO_3_ as N source (Table 1). *Mtnr3* was not analysed in this experiment as it is only expressed during symbiotic interaction. As a control, "sibling" lines (*nr1_Sibl1* and *nr1_Sibl2* described in Table 2) were used to confirm that the observed phenotypes are not associated with *Tnt1* insertions unrelated to the NR genes of interest that may have persisted in the lines after several backcrosses. NR activity was measured in both leaves and roots of the *Mtnr1* and *Mtnr2* mutant compared to WT. The results showed a 90% and 70% reduction in NR activity in shoots in the *Mtnr1* and *Mtnr2* mutants respectively (Table 1). It is noteworthy that the combined effect of *Mtnr1* and *Mtnr2* exceeded 100%. In the *Mtnr1/nr2* double mutant, the NR activity was only 3% in aerial parts and 2% in roots. As a control, the *nr1_Sibl1* line exhibited a 90% decrease in NR activity, which was similar to the *Mtnr1* line. These results confirm that there is no phenotype caused by potential residual *Tnt1* mutations in the analyzed lines. Altogether, these data confirm the major impact of the inactivation of NR1 and NR2 on the NR activity of nitrate-fed plants. In the same experiment, a significant 20% reduction in shoot and root biomass was observed in the *Mtnr1* mutant, the *nr1_Sibl1* line and the *nr1_Sibl2* line compared to the WT control (Fig. 3A and B), whereas the *nr2* mutant was not affected, indicating a major effect of the NR1 isoform on nitrate assimilation by the plant. In the double mutant, root biomass was reduced by 60 % and shoot biomass by 92 % compared to the WT (Fig. 3A and B). Consequently, the root/shoot ratios significantly increased in the double mutant, while no noticeable variation was observed in the other lines (Fig. 3C). This phenotype was already observed under severe nitrogen deficiency conditions (Lopez et al., 2023). The production of NO was analyzed in the root of the double mutant lines and control grown on 2 mM nitrate. Interestingly, we did not observe any reduction in the production of NO by the roots of the double NR mutant (Fig. 4) suggesting that the production of NO, as measured on non-inoculated roots, is not dependent on NR activity.

**Figure 3:**
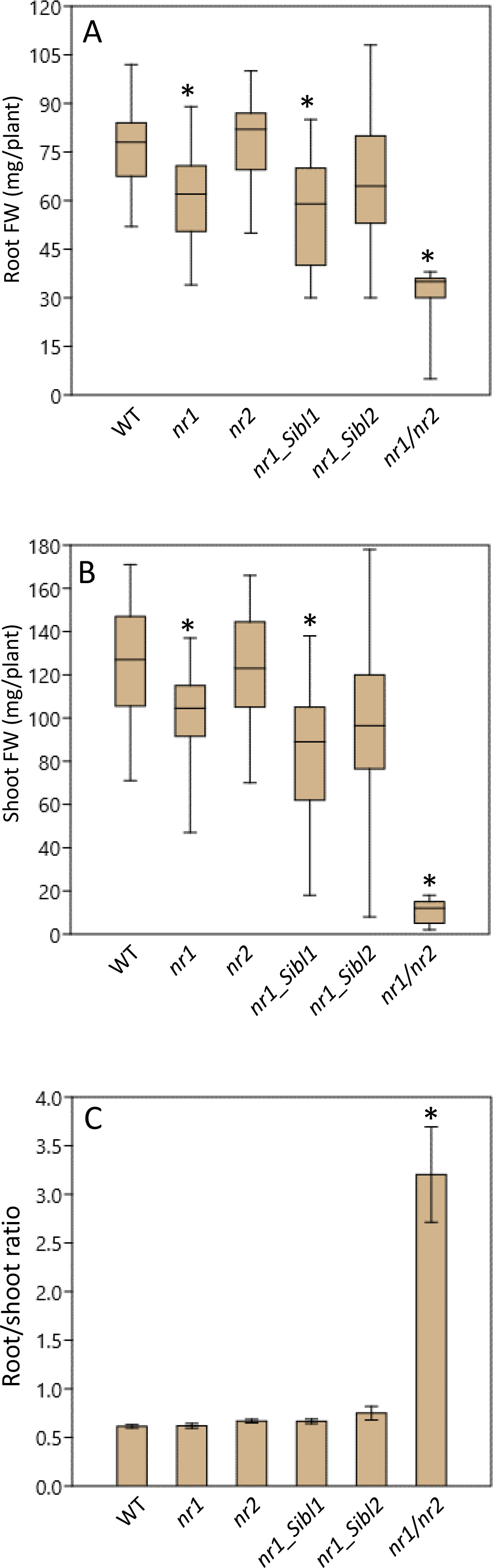
**Biomass measurements of different NR mutants growing on nitrate** (A) Measurement of root biomass after 2 weeks of growth on NO_3_^-^. (B) Measurement of shoot biomass after 2 weeks of growth on NO_3_^-^. (C) Root/shoot ratio measurement. The asterisks correspond to significant differences determined by the Mann–Whitney test (p < 0.05) from two independent experiments composed of, at least, 25 plants for each condition.

**Figure 4:**
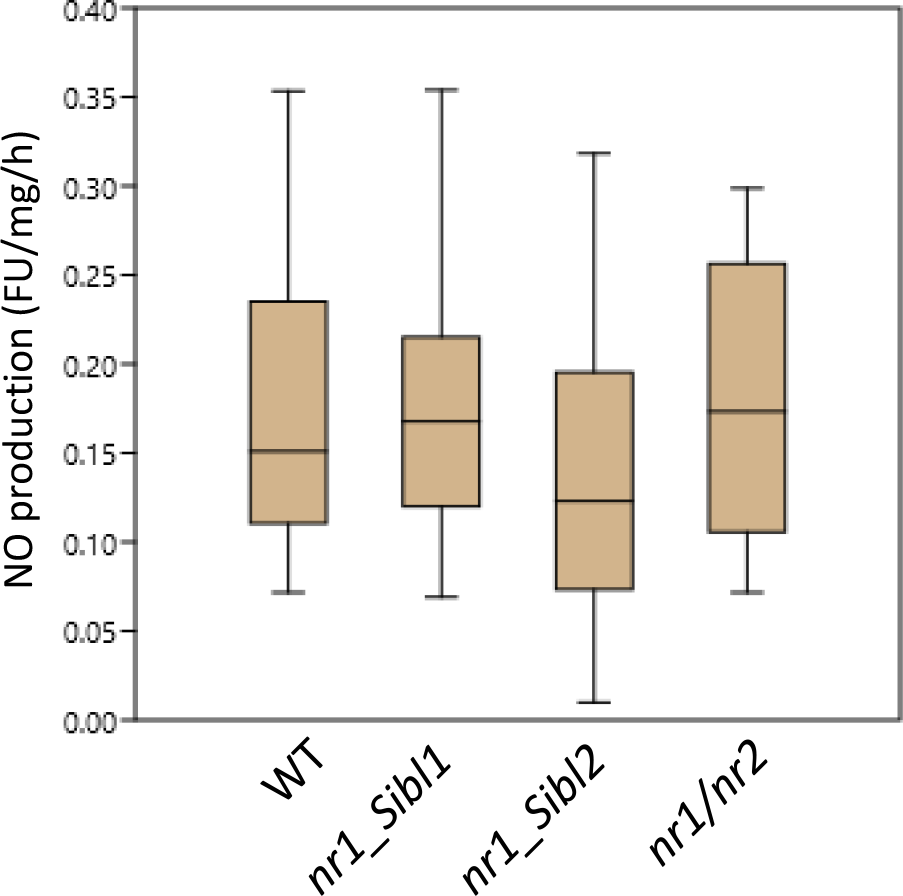
**Measurement of NO production from non-inoculated root**. The fluorescence intensity of the NO production was measured using the 4,5-diaminofluorescein probe (DAF-2;Sigma-Aldrich). The line within the boxes refers to the median. The asterisks correspond to significant differences determined by the Mann–Whitney test (p < 0.05) from two independent experiments composed of, at least, 12 plants for each condition. Each measurement was done in triplicate.

**Table 1:** Measurement of nitrate reductase activity in shoots, roots and nodules of *M. truncatula* NR mutants. The nitrate reductase activity is expressed in nmol per g of fresh weight per h. . Results are the mean ± SE (*p < 0.05, one-way analysis of variance) of two independent experiments composed of, at least, 25 plants for each condition. Each measurement was done in triplicate.

|  |  | WT | *nr1* | *nr2* | *nr3* | *nr1_Sibl1<sup>o</sup>* | *nr1/nr2* |
| --- | --- | --- | --- | --- | --- | --- | --- |
| NR activity<br>(nmol/g of FW/h) | Leaves | 1084 ± 258 | 123 ± 30 | 306 ± 76 | | 112 ± 14 | 31 ± 12 |
| | Roots | 298 ± 95 | 27 ± 10 | 110 ± 20 | | 57 ± 19 | 5.2 ± 3 |
| | Nodules | 229 ± 62 | 39 ± 21 | 116 ± 44 | 224 ± 56 | | |

**Table 2:** Nomenclature of mutant lines used in this manuscript.

| Name | genotype | <sup>o</sup> |
| --- | --- | --- |
| *nr1* | *nr1-1<sup>ko</sup>* | KO for the *NR1* gene |
| *nr2<sup>o</sup>* | *nr2-1<sup>ko</sup>* | KO for the *NR2* gene |
| *nr3* | *nr3-1<sup>ko</sup>* | KO for the *NR3* gene |
| *nr1_Sibl1* | *nr1<sup>ko</sup>* | KO for the *NR1* gene with no *Tnt1* insertion in the *NR2* gene |
| *nr1_Sibl2* | *nr1<sup>ko</sup>_nr2<sup>HT</sup>* | KO for the *NR1* gene and heterozygous on *NR2* gene allele |
| *nr1/nr2* | *nr1/nr2-1<sup>ko</sup>* | KO for the *NR1* and *NR2* gene |

#### Impact of *M. Medicago* NR mutants on the nodulation process

In previous studies, it was shown that NR activity is required for the N_2_-fixing symbiosis (Horchani et al., 2011; Berger et al., 2020a). Three simultaneous peaks of *MtNRs* expression and NR activity were observed during the first hours of symbiotic interaction at 10 hours post-inoculation (hpi), during early nodule development at 4 days post-inoculation (dpi), and when the nodule matures around 3-4 weeks post-inoculation (wpi) (Berger et al., 2020a). To investigate the role of each isoforms during the nodulation process, the three single *nr* mutants were grown in symbiosis with *S. meliloti*. The measurement of NR activity shows that KO of the *NR1*, *NR2*, or *NR3* genes resulted in an 87%, 50%, and 3% decrease in activity in 4-week-old nodules, respectively, as shown in Table 1. However, no significant difference between mutants and WT was found for nodule number, plant biomass (shoot dry weight), and nitrogenase activity (acetylene reduction assay, Fig S4). Considering that NR was shown to be a source of NO during the establishment and functioning of N_2_-fixing symbiosis (Berger et al., 2020a), we investigated the participation of each NR in NO production in mature nodule. The NO production in 4-week-old nodules was not affected for the 3 *Mtnr* mutants (Fig. 5). The NO level in the nodules of *M. truncatula* depends not only on the enzymes involved in its synthesis, but also on those involved in its catabolism such as the Phytoglobine1.1 (Berger et al., 2018). The analysis of *MtPgb1.1* expression in 4 wpi nodules showed that *Pgb1.1* was not significantly increased in the different mutant (Fig. 6). Since *Pgb1.1* is regulated by NO at the transcriptional level (Berger et al., 2020b), the lack of regulation in its expression is consistent with the absence of variation in NO levels in the three mutants. In addition, qPCR analysis revealed that the absence of one of the three NRs in the three mutants did not result in overexpression of the other genes to compensate (Fig. 6). Expression of *MtNiR* gene, known to be overexpressed in *nr* mutant of *Nicotiana plumbaginifolia* (Faure et al., 1991), was not affected in the three simple *nr* mutants. The absence of phenotype observed in single mutants suggests significant functional redundancy between the different NRs in *M. truncatula* nodules.

**Figure 5:**
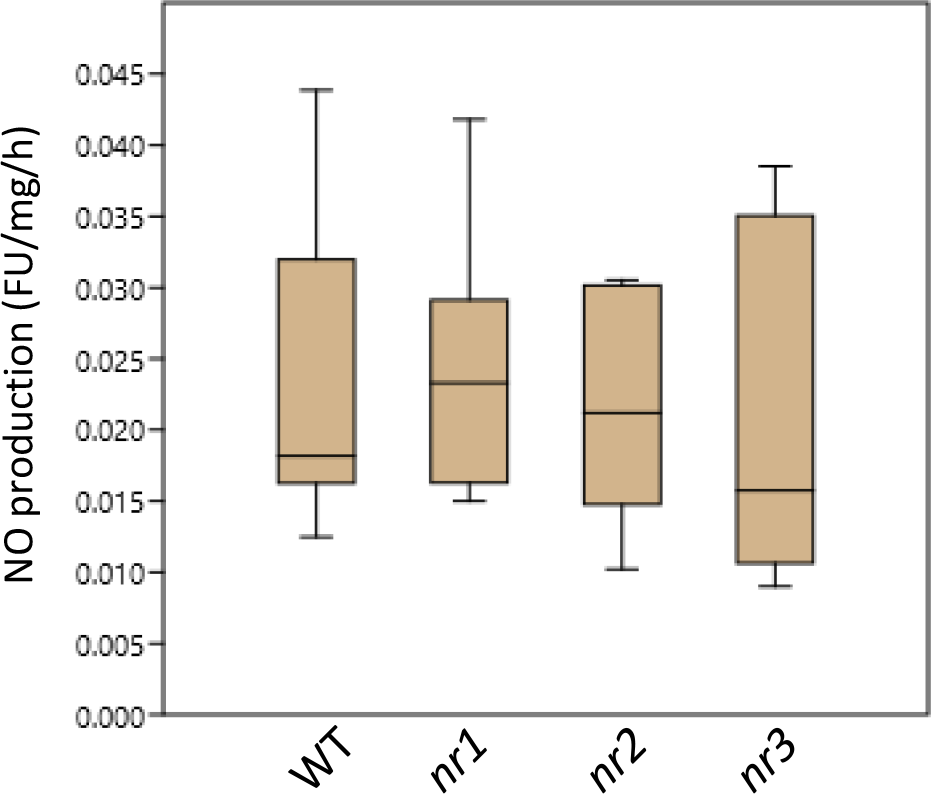
**Measurement of NO production from mature nodule**. The fluorescence intensity of the NO production was measured using the 4,5-diaminofluorescein probe (DAF-2;Sigma-Aldrich). The line within the boxes refers to the median. The asterisks correspond to significant differences determined by the Mann–Whitney test (p < 0.05) from two independent experiments composed of, at least, 12 plants for each condition. Each measurement was done in triplicate.

**Figure 6:**
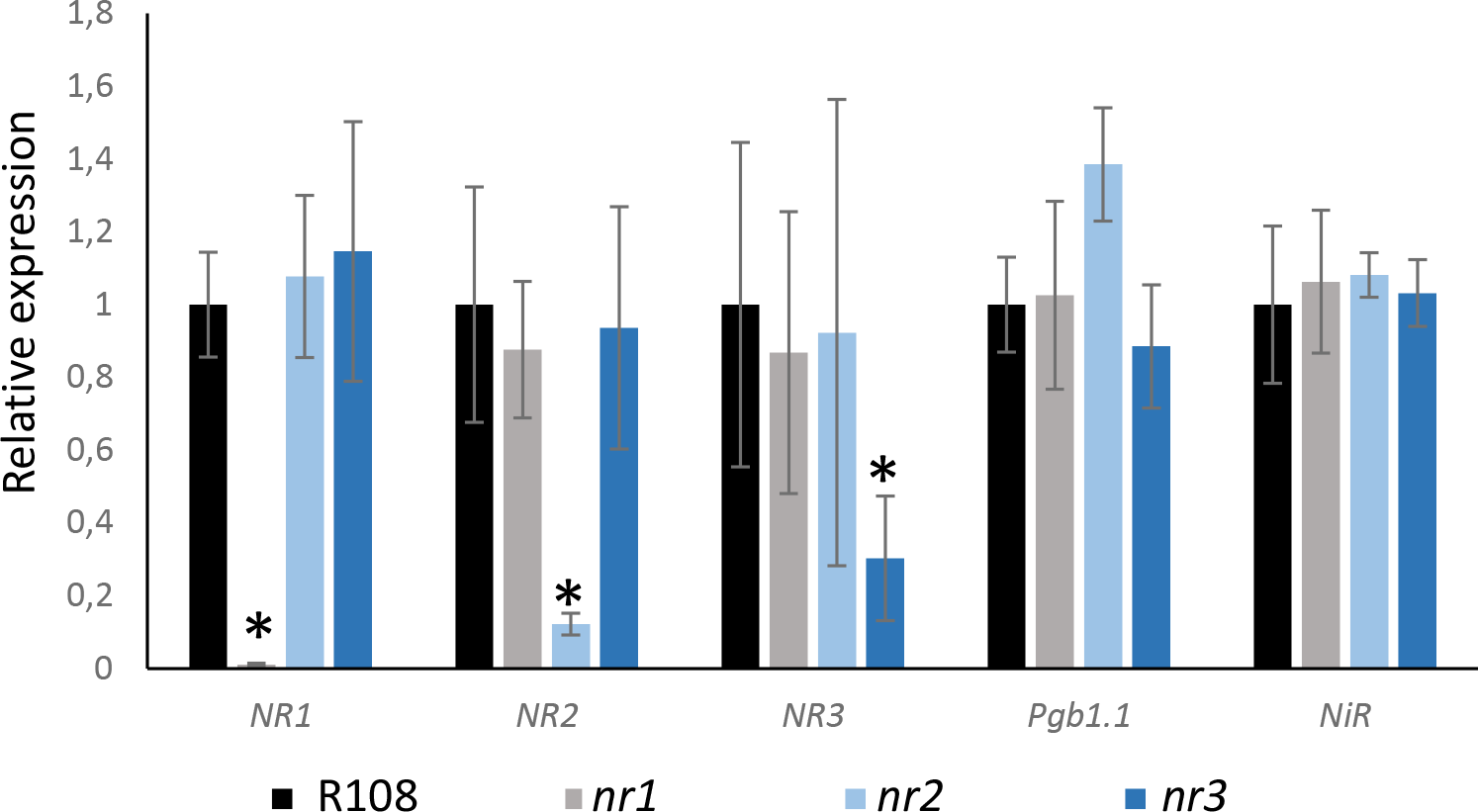
**Expression analysis of the three *NRs, Pgb1.1 and Nir* genes in nodules from NR mutants.** Relative transcript levels were measured by real-time quantitative PCR on 4 week-old *M. truncatula* nodules, from 4 different genotypes (R108, *Mtnr1*, *Mtnr2* and *Mtnr3*). Results are the mean ± SE. The asterisks correspond to significant differences determined by the Mann–Whitney test (p < 0.05) from two independent experiments composed of, at least, 10 plants for each condition.

We therefore analyzed the effects of the double mutation, *NR1* and *NR2*, on the nodulation process. Hydroponic cultures were performed on segregating mutant lines, *nr1/nr2* double mutant, *nr1_Sibl1*, and *nr1_Sibl2* lines inoculated with *S. meliloti*. Figure 7 shows that only the *nr1/nr2* double mutant was affected in terms of nodule number and root biomass. The double mutant showed a 85% reduction of nodule number at 7 dpi, reaching 98% reduction at 4 wpi (Fig. 7B and D). Regarding the growth, at 7 days post-inoculation, the double mutant already displayed a 50% decrease in root DW biomass in comparison to the control (Fig. 7A), while the shoot DW decreased by only 25% (Fig S5A). At 4 wpi, the biomass of both the shoots and the roots was reduced by 70% in the double mutant as compared to all the other lines analysed (Fig. 7B and Fig. S5B). In view of the almost complete absence of mature nodules in the double mutant, it was not possible to analyse the production of NO in nodules devoid of NR activity.

**Figure 7:**
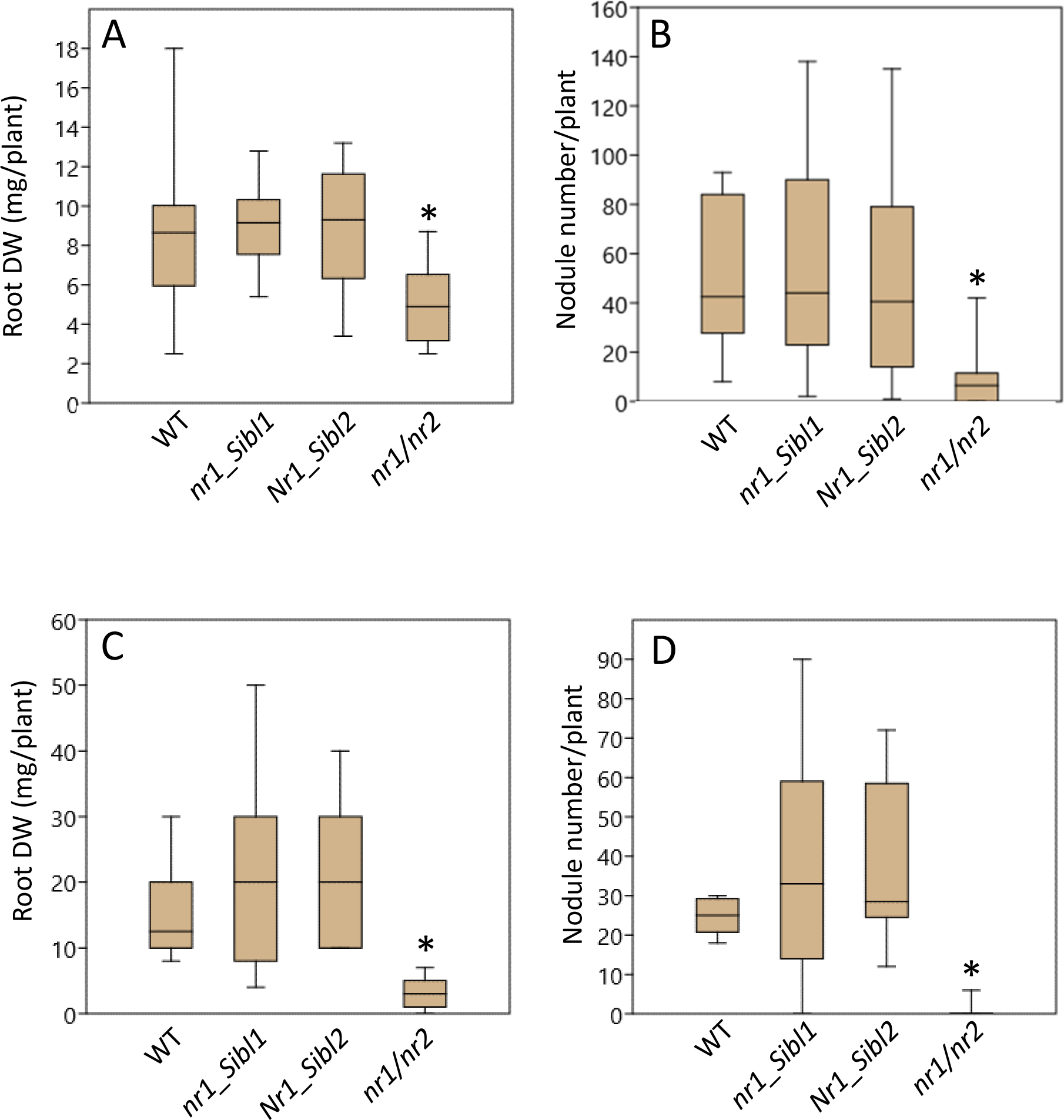
**Phenotype of *M. Medicago* NR mutants during the nodulation process.** Boxplot representation of root dry weight and nodule number measured on hydroponic culture of segregating double mutant after 7 days (A and B) and 4 weeks (C and D) post inoculation with *S. meliloti*. The line within the boxes refers to the median. The asterisks correspond to significant differences determined by the Mann–Whitney test (p < 0.05) from two independent experiments composed of, at least, 15 plants for each condition.

NR activity was previously shown to be involved in nodule energy metabolism via the "Phytogb-NO" respiration (Horchani et al., 2011; Berger et al., 2021). Thus, the reduced NR activity in *nr1* and *nr2* mutants should potentially lead to a reduced energy metabolism and a large metabolic reprogramming even in a single mutant. Therefore , we investigated the effect of NR knock-out on the nodule metabolome. After GC/MS analysis, 202 chemical species were detected in the nodules among which 102 were identified (Table S3).. Most of them were soluble sugars, amino acids, and organic acids (Table S3). The results showed that there was no major difference in the nodule metabolome of the 3 mutants as compared to the WT, excepted for serine, 2-piperidinecarboxylate, citrate and glutaric acid (Fig. 10). The absence of contrasting phenotype for metabolite content between the mutants and the WT suggests that even a 10% residual NR activity is apparently sufficient to maintain the nodule metabolism.

## Discussion

Numerous investigations were conducted on nitrate reductases from 1970 to 2000, elucidating the structure, function and regulation of nitrate reductase from higher plants (reviewed in (Campbell, 1999)). However, despite the identification of NR mutants in legumes (Feenstra and Jacobsen, 1980; Ryan et al., 1983), no study has fully elucidated the significance of these enzymes in the process of nitrogen-fixing symbiosis. The role of NR together with that of NiR in the symbiotic relationship between legumes and rhizobia is notably questioned because atmospheric nitrogen (N_2_) is directly reduced to ammonia (NH_3_) by bacterial nitrogenase. Since 2000, there has been a renewed interest in NR due to its role in NO synthesis in plants (Dean and Harper, 1988; Yamasaki and Sakihama, 2000; Rockel et al., 2002; Sakihama et al., 2002; Lea et al., 2004; Gupta et al., 2005; Planchet et al., 2005; Kolbert et al., 2010). More recent reports demonstrated that NR plays a role in generating nitrite and indirectly nitric oxide (NO) during the establishment and functioning of the N_2_ fixation symbiosis (Horchani et al., 2011; Berger et al., 2020a). Three genes encoding NRs, namely *MtNR1*, *MtNR2*, and *MtNR3*, were reported in the *M. truncatula* genome (Puppo et al., 2013). The availability of invalidated mutants for the three NRs of *M. truncatula* provides an opportunity to unveil the distinctive roles played by each of these NRs during the nodulation process.

### Phylogenetic analysis of the NT protein family

NRs are widespread proteins in plants playing a crucial role in nitrogen metabolism, making them exceptional indicators for monitoring evolutionary processes over extended periods (Opperdoes and Lemey, 2009). Phylogeneticy analysis of the NR family in the Rosids clade was facilitated by the recent increase in sequenced genomes (Fig. 1, Table S2). The results showed that a significant number of species in the Rosids clade possess two NR sequences. Each order belonging to the Rosids clade, except for Myrtales, forms a monophyletic group. This suggests that duplication events responsible for the existence of different *NR* genes, occurred after the divergence of the different orders within the Rosids clade. This statement implies that the different NR sequences in Fabales and Brassicales orders may have diverged originated from independent duplication events. This means that a direct comparison of the function of NRs from *M. truncatula* with those from *A. thaliana* should be made with caution. The phylogenetic analysis indicates that a third NR gene, named NR3-type in this work, has emerged from a duplication of *NR1* in the inverted repeat-lacking clade (IRLC) of the Fabales order (Fig. 1). In *M. truncatula*, *MtNR1* and *MtNR3* are found to be separated by only 20 kb on the same chromosome (Berger et al., 2020a) suggesting that the duplication of *NR1* to give rise to *NR3* occurred by unequal crossing-over. Although three NRs sequences were identified in *Trifolium subterraneum* and *M. truncatula*, only two sequences were found in the genomes of the other three species in the IRLC, *Pisum sativum*, *Vicia faba* and *Cicer arietinum*. Interestingly, these three species appear to have retained the *NR3*-type gene while losing the *NR1* gene sequence (Fig. 1). IRLCs form nodules of indeterminate type and have the peculiarity of expressing NRC (Nodule Cysteine-Rich) peptides, which are specific to nodules and play a key role in terminal bacteroid differentiation (Czernic et al., 2015). More investigations are needed to discern whether the emergence of a third NR in IRLC is indicative of a neofunctionalization event. It would therefore be interesting to elucidate the potential roles of these NR3-type NRs in indeterminate nodules and whether this enzyme is part of a specific control mechanism that has emerged in some legumes of the IRLC clade.

The alignment of all the identified NR sequences unveiled a new intriguing feature in the NR3-type sequences of 5 Fabales species. These five NR proteins exhibit differences in the two hinge regions that, are typically highly conserved across species. Previous studies have demonstrated that phosphorylation of Serine 534 in *A. thaliana* (Fig. 2B), located in Hinge 1 between the Moco and heme domains, results in the formation of a complex between NR and 14-3-3 proteins (Lea et al., 2004; Lillo et al., 2004). This renders NR more susceptible to proteome-mediated degradation, as reported by Lillo et al. (2004). Additionally phosphorylation of Serine 627 in Hinge 2 of *A. thaliana* has been observed as a rapid activation mechanism in response to the need for excessive nitrate reduction (Wang et al., 2010). The results suggest that this type of NR may employ a different regulatory mechanism compared to other NRs.

Furthermore, Berger and colleagues (2020a) proposed that MtNR3, exclusively expressed during symbiotic interaction, may play a role in the onset of nodule senescence. Additionally, the authors demonstrated that the presence of nitrate inhibits *NR3* expression, while inducing that of *NR1* and *NR2* (Berger et al., 2020a). Discrepancies in NR3 sequence as compared to NR1 and NR2 may result in lack of regulation by phosphorylation, thus conferring increased resistance to proteolysis (Lillo et al., 2003; Lambeck et al., 2012), a process occurring during nodule senescence approximatively 6-8 weeks post-inoculation with *S. meliloti*. This resistance could allow NR3 to participate in the nodule senescence process, as observed by Berger et al. (2020a). However, nitrogen fixation in mature nodules was not affected in the *Mtnr3* mutant line (Fig. S3C). Other analyses, such as growth (Fig. 3), NO production (Fig. ) or metabolism (Fig. 6), performed on the *Mtnr3* mutant failed to identify a role for NR3 in the establishment of symbiotic interaction up to the onset of nodule senescence. One possible explanation for the lack of phenotype may be due to its highly specific expression profile in the nodule meristem and its lower expression level compared to the other two NRs in the nodule, an assumption supported by an absence of NR activity reduction in the *Mtnr3* mutant roots and nodules (Table 1). Given that *MtNR3* from *M. truncatula* is exclusively expressed during symbiotic interaction (Berger et al., 2020a), we investigated if the other *NR3-*type *NRs* from IRLC exhibited a similar expression pattern than *MtNR3*. Database available for *P. sativum* and *C. arietanum* showed that *PsNR3* and *CaNR3* are well expressed in different organs of the plant and not exclusively to the nodule. Therefore, it appears that the presence of the third NR branch in IRLC is not necessarily correlated with the nodulation process.

Additionally, it is noteworthy that a third NR sequence emerges within the *Cucurbitales* order. Upon aligning the three NR sequences of *C. sativus* with other NR sequences (Fig. 2), it becomes apparent that phosphorylation site is lost in NR3-type sequence (Cusat_NP_001267688), similar to the IRLC. The *Cucurbitales* order has also exhibited this specificity, despite its inability to engage in symbiotic interaction. This suggests that the absence of conserved phosphorylation sites is not directly associated with the functioning of undetermined type nodules in IRLC. It should be noted that a transgenic *N. plumbaginifolia* line expressing a mutated tobacco NR, where Ser 521 has been changed to Asp by site-directed mutagenesis, was shown to have a permanently active NR enzyme (Lillo et al., 2003). This mutation completely abolishes inactivation in response to light/dark transitions or other treatments that are known to inactivate NR. Indeed, in wild-type plants, NR is in the inactivated form during prolonged darkness, whereas in the S521 line, NR is always in the active form (Lillo et al., 2003). In the S532D and S532A mutants of the *NIA1* gene in *Oryza sativa*, the level of phosphorylation was reduced, leading to an improvement in nitrate assimilation but a reduction in plant height (Han et al., 2022). The authors suggested that an increase in phosphorylation levels reduces the accumulation of nitrite and prevents the toxic effects of reactive oxygen species in rice (Han et al., 2022). The appearance of this type of NR3, without phosphorylation sites, may therefore have been guided by an advantage of this always active form in IRLC as in *Cucurbitales* order not necessary with the same objective.

### Phenotype of the Medicago NRs mutants

Overall, the absence of phenotype observed in *M. truncatula* single mutants suggests significant functional redundancy between the different NRs in *M. truncatula*. The most striking outcome of this work is the effect of the absence of *NR1* and *NR2* on nodulation efficiency. Despite the presence of MtNR3, the double mutant is strongly affected on nodulation with 85% reduction of nodule number at 7 dpi, reaching 98% reduction at 4 wpi. A result that demonstrates that NR activities are required for the functioning of the N_2_-fixing symbiosis.

A previous study showed that *MtNR2* is the most highly expressed *NR* gene (30 to 100 times more than *NR1*) throughout the whole process in *M. truncatula* (Berger et al., 2020a). Surprisingly, the reduction in NR activity is more pronounced in the *Mtnr1* mutant (90%) than in the *Mtnr2* (70%) or *Mtnr3* (less than 1%) mutants. The *Mtnr1* mutant retains only 10% of the WT shoot NR activity as observed for the *NIA2* mutant of Arabidopsis (Wilkinson and Crawford, 1991). In contrast to the *A. thaliana NIA2* mutant, which grows normally with nitrate as the sole nitrogen source, the *Mtnr1* mutant exhibited a significant 20% reduction in shoot and root biomass compared to the WT control (Figure 4A and B). On the other hand, the growth on nitrate of the *Mtnr2* mutant which possess again 30% of the WT shoot NR activity was not affected (Table 1). These results are in favour of a major role of the NR1 isoform on nitrate assimilation by the plant. It is noteworthy that the combined effect of *Mtnr1* and *Mtnr2* exceeded 100% in leaves, roots and nodules. These results indicate that both NR1 and NR2 must be present to obtain full NR activity, indicating a potential cooperative effect in total NR activity. In the *Mtnr1/nr2* double mutant, the NR activity was only 3% in aerial parts and 2% in roots compared to the control. These results are closed to the 0.5% of wild-type shoot NR activity observed in *Atnia1/nia2* double mutant (G′4-3 mutant) of Arabidopsis (Wilkinson and Crawford, 1993) and to the total absence of NR activity in the NR-full mutant (Wang et al., 2004). Unlike the Arabidopsis G′4-3 double mutant, which shows very poor growth on media with nitrate, the Medicago double mutant *Mtnr1/nr2*, as a Arabidopsis NR-full mutant, does not grow under the same conditions. Furthermore, contrary to Arabidopsis NR-full mutant and in spite of numerous attempts of growth experiments with different sources of nitrogen, it was not possible to get to the point of flowering and seed production in the double mutant seedlings that are knocked out (KO) on the *NR1* and *NR2* genes.

Nitrate reductase (NR) activity is typically present in legume root nodules of *Lotus japonicus* (Kato et al., 2003), *Glycine max* (Kanayama et al., 1999) and *M. truncatula* (Horchani et al., 2011; Berger et al., 2020a). The localization of NR mRNA within the infected regions of pea root nodules (Kato et al., 2003) and *M. truncatula* nodule (Horchani et al., 2011) suggest a specific function in this organ. This is supported by the fact that NR activity is required for N_2_ fixing symbiosis and nodule energy metabolism (Horchani et al., 2011; Aridhi et al., 2020; Chammakhi et al., 2022). Furthermore, recent study showed a significant increase in the expression of *MtNR1* and *MtNR2* at 10 hpi, 4 dpi, and 5 wpi, which coincides with a rise in NO levels (Berger et al., 2020a). *MtNR1*, *MtNR2* and *MtNR1-NR2* Knock-down using a nodule-targeted RNA interference (RNAi) strategy led to a decrease by about 50% of NR activity in the three RNAi constructions with a concomitant reduction of NO production (Berger et al., 2020a). Furthermore, nodule size on *MtNR1/2 RNAi* transgenic roots was reduced by 30% to 40% (Horchani et al., 2011). In contrast to the results obtained by RNAi, the study of the different NR KO failed to reveal the contribution of any of these NRs to the establishment and/or functioning of the nodule even if the KO mutation of *Mtnr1*, *Mtnr2* and *Mtnr3* resulted in 87%, 50% and 3% reduction in NR activity compared to control. In the same way, none of these 3 mutants showed a decrease in NO production (Fig. 6). These results suggest that the NR expressed in *M. truncatula* are complementary and that only 13% NR activity is sufficient to maintain the proper functioning of the nodule.

In nodules derived from the *Mtnr3* mutant, NO production remains unaffected. A result that could be due to minor role of NR3 in total NR activity or an absence of phosphorylation in hinge 2. Indeed, previous research on *A. thaliana* has indicated that phosphorylation at serine 627 of the Nia2 isoform results in a notable rise in its activity and NO production (Wang et al., 2010). Likewise, the overexpression of *NIA1* and, to a lesser extent *NIA2*, led to increasing NO accumulation, thus suggesting that specific NR are involved in NO in Arabidopsis (Costa-Broseta et al., 2021). And although previous work with RNAi on MtNR suggested a preferential involvement of MtNR1 in NO production (Berger et al., 2020a), NO production was not affected in each single mutant in roots and nodules. The mechanism by which NO is synthesized in plants remains controversial and incompletely understood. It is important to note that NR is not the sole source of NO synthesis in plants (Berger et al., 2018; Kolbert et al., 2019). Additionally, it should be kept in mind that NO production in the nodule is not solely attributed to the plant partner (Horchani et al., 2011; Salas et al., 2021). Research has shown that in *M. truncatula* and soybean nodules, the bacterial partner produces between 33% to 90% of NO (Sánchez et al., 2010; Horchani et al., 2011). We can therefore postulate that in NRs mutants, nitrate is reduced to nitrite mainly in bacteroids. This would explain why NO production did not vary.

NR was suggested to play a role in the metabolism, in particular the energy metabolism in the nodule of *M. truncatula* (Berger et al., 2020a). Previous results demonstrate that inhibition of NR by tungstate (Tg), with or without NO_3_^-^, lead to an increase in sucrose content, indicating a decrease in its consumption by the nodules and resulting in lower levels of succinate and malate (Berger et al., 2020a). Metabolomic analyses in our work did not reveal any differences in malate, succinate or sucrose content in 4 weeks old nodules for the 3 single mutants. It should be noted that Berger et al (2020a) investigations have been done on detached nodules treated with tungstate for 4 h. It is conceivable that 4 h treatment with Tg leads to transient metabolic disturbances that disappear in *Mtnr* mutants as a result of their long term adaptation to low level of NR activity.

In conclusion, an important point of this work is the identification of a third NR sequence, here called NR3-type, which arose in the inverted repeat-deficient clade of the Fabales order. An intriguing feature of the NR3-type sequences is the absence of conserved phosphorylation sites in the two hinge regions, in contrast to all other NRs. The appearance of this type of NR3, without phosphorylation sites, may therefore have been guided by an advantage of this always active form in IRLC. However, *M. truncatula* MtNR3, which is only expressed in nodules, does not seem to play an important role. Our results showed that *M. truncatula* NRs are complementary and perform the same function in nitrogen metabolism even though MtNR1 predominates in the total NR activity measured in shoots, roots and nodules. Then, contrary to Arabidopsis NR1 and NR2, none of the 3 Medicago NRs was found to play a more specific role in NO production (Wang et al., 2010). This conclusion is consistent with previous work describing NRs that are not directly involved in NO synthesis in *M. truncatula* (Horchani et al., 2011; Berger et al., 2020a; Berger et al., 2021). Furthermore, only 13% NR activity is sufficient to maintain the proper functioning of the nodule. Interestingly, the double mutant line *Mtnr1/nr2* conducted to an almost complete absence of nodulation therefore demonstrating for the first time the importance of NR in N_2_-fixing symbiosis. Unfortunately, this absence of nodules makes it difficult to pinpoint the precise role of NR activity in nodule functioning. Previous work of our team showed that NR activity play a key role in nodule energy metabolism via the "Phytogb-NO" respiration (Horchani et al., 2011; Berger et al., 2020a; Berger et al., 2020b). Hence, the absence of nodulation in this NR-deficient double mutant reinforces the major role of this “Phytogb-NO” respiration in nodule functioning. This work also raises also the question of the involvement of the bacteroid in the maintenance of NO homeostasis in the nodule, which is necessary for its proper functioning.

## Author Contributions

Conceptualization, BA, BR, LM; Investigation BM, DA, HAN, CD, PM, EFY, LM, CG, BR, BA; Roles/Writing – original draft, BA, HAN, BR; Writing -review & editing BA, BR, LM, CD, HAN; Supervision, BA. All authors have read and agreed to the published version of the manuscript.

## Supplemental materials

Table S1 : List of primers used in the manuscript for cloning and quantitative RT-PCR analysis.

Table S2: **Table of all NR sequences data** found from an exhaustive search of 3 genomic and protein databases: Phytozome (www.phytozome.jgi.doe.gov), Uniprot (www.uniprot.org) and NCBI (www.blast.ncbi.nlm.nih.gov). The NR sequences have been named by a code composed of 5 letters and numbers representing the species to which they belong, the name that may have been assigned to them from the literature and their amino acid number.

Table S3: Mature nodule metabolome of the 3 mutants as compared to the WT.

Figure S1 : **Maximum likelihood phylogenetic tree of NR protein sequences belonging to 10 different orders of the Rosids clade.** NR sequences from the Poales clade were used as an outgroup to root the phylogenetic tree. The phylogeny was constructed from 121 different NR protein sequences, ranging from 582 to 956 amino acids, from 87 species (Table S1). Red dots on selected nodes indicate the duplication events at the origin of the well-characterized NIA1 and NIA2 from *A. thaliana* and NR1, NR2, and NR3 from *M. truncatula* (indicated by a green dot outside the phylogeny). Black dots indicate nodes with support values greater than or equal to 80% and 95% for SH-aLRT and UFboot, respectively.

Figure S2. **Expression of *C. arietanum* and *P. sativum NRs* genes in different part of each plant.** (A) The RNA-seq data obtained for the various reproductive tissues analysed by Kudapa et al. (2018) are expressed in raw FPKM. Following phylogenetic results, *Ciari_XP004513831* gene was named *CaNia2* and *Ciari_XP004500600* was named *CaNR3*. (B) Expression profiles of the pea *NR* gene expressed in RPKMnormalised were retrieved from the Pea Gene Atlas (Alves-Carvalho et al., 2015). RPKMnorm is the RPKM divided by the geometric mean of the RPKM for three control genes: histone H1 (*PsCam009820*), actin (*PsCam042218*) and EF1a (*PsCam042119*). Stage A represents 7–8 nodes, 5–6 opened leaves; stage B represents the start of flowering; stage C represents 20 days after the start of flowering Following phylogenetic results, *Pisat_884* gene was named *PsNR2* and *Pisat_869* gene was named *PsNR3*.

Figure S3: **Identification of *Tnt1*-insertion line NF15469, NF14768 and NF11190 for each *NR* gene of *M. truncatula*.** A. Schematic representations of *Tnt1* retrotransposon localisation in *MtNR1*, *MtNR2* and *MtNR3* genes respectively as determined by PCR reverse screening.

Exons are depicted as black boxes, and introns and untranslated regions are represented by lines. Scale is 1 kb only the Tnt1 retrotransposon is not to scale on the representation.

Figure S4: Nodule number, plant biomass and nitrogen-fixing capacity of the different ***M. truncatula* NR mutants:** (A) Nodule number was determined on plants growing in plate with Fahraeus medium at 14 days post inoculation. (B) Plant biomass was measured on the shoot of the plant used for acetylene reduction assay (ARA). (C) The nitrogen-fixing capacity was measured by the ARA expressed in nmol of ethylene (C_2_H_4_) reduced per mg of nodules and per hour, from plant nodules at 4 weeks post inoculation (wpi). The asterisks correspond to significant differences determined by the Mann–Whitney test (p < 0.05) from two independent experiments composed of, at least, 9 plants for each condition.

Figure S5: **Plant biomass of *M. truncatula* NR double mutants during the nodulation process.** Boxplot representation of shoot DW measured after 7 days (A) and 4 weeks post inoculation with *S. meliloti* on hydroponic culture of segregating double mutant. The line within the boxes refers to the median. The asterisks correspond to significant differences determined by the Mann–Whitney test (p < 0.05) from two independent experiments composed of, at least, 15 plants for each condition.

## References

1. Aridhi F, Sghaier H, Gaitanaros A, Khadri A, Aschi-Smiti S, Brouquisse R (2020) Nitric oxide production is involved in maintaining energy state in Alfalfa (Medicago sativa L.) nodulated roots under both salinity and flooding. Planta 252: 22

2. Arrese-Igor C, García-Plazaola JI, Hernández A, Aparicio-Tejo PM (1990) Effect of low nitrate supply to nodulated lucerne on time course of activites of enzymes involved in inorganic nitrogen metabolism. Physiol Plant 80: 185–190

3. Bachmann M, Shiraishi N, Campbell WH, Yoo BC, Harmon AC, Huber SC (1996) Identification of Ser-543 as the major regulatory phosphorylation site in spinach leaf nitrate reductase. Plant Cell 8: 505–517

4. Berger A, Boscari A, Horta Araújo N, Maucourt M, Hanchi M, Bernillon S, Rolin D, Puppo A, Brouquisse R (2020a) Plant Nitrate Reductases Regulate Nitric Oxide Production and Nitrogen-Fixing Metabolism During the Medicago truncatula–Sinorhizobium meliloti Symbiosis. Front. Plant Sci. 11:

5. Berger A, Boscari A, Puppo A, Brouquisse R (2021) Nitrate reductases and hemoglobins control nitrogen-fixing symbiosis by regulating nitric oxide accumulation. J Exp Bot 72: 873– 884

6. Berger A, Brouquisse R, Pathak PK, Hichri I, Inderjit, Bhatia S, Boscari A, Igamberdiev AU, Gupta KJ (2018) Pathways of nitric oxide metabolism and operation of phytoglobins in legume nodules: Missing links and future directions. Plant Cell Environ 41: 2057–2068

7. Berger A, Guinand S, Boscari A, Puppo A, Brouquisse R (2020b) Medicago truncatula Phytoglobin 1.1 controls symbiotic nodulation and nitrogen fixation via the regulation of nitric oxide concentration. New Phytol 227: 84–98

8. Bosseno M, Lambert A, Beucher D, Le Gleuher M, Aubry C, Pauly N, Montrichard F, Boscari A (2020) A simple method for genetic crossing in Medicago truncatula. Model Legume Medicago Truncatula. John Wiley & Sons, Ltd, pp 1027–1033

9. Campbell WH (2001) Structure and function of eukaryotic NAD(P)H:Nitrate reductase. Cell Mol Life Sci 58: 194–204

10. Campbell WH (1988) Nitrate reductase and its role in nitrate assimilation in plants. Physiol Plant 74: 214–219

11. Campbell WH (1999) Nitrate reductase structure, function and regulation: Bridging the gap between biochemistry and physiology. Annu Rev Plant Biol 50: 277–303

12. Capella-Gutiérrez S, Silla-Martínez JM, Gabaldón T (2009) trimAl: a tool for automated alignment trimming in large-scale phylogenetic analyses. Bioinformatics 25: 1972–1973

13. Chammakhi C, Boscari A, Pacoud M, Aubert G, Mhadhbi H, Brouquisse R (2022) Nitric Oxide Metabolic Pathway in Drought-Stressed Nodules of Faba Bean (Vicia faba L.). Int J Mol Sci 23: 13057

14. Clément G, Moison M, Soulay F, Reisdorf-Cren M, Masclaux-Daubresse C (2018) Metabolomics of laminae and midvein during leaf senescence and source-sink metabolite management in Brassica napus L. leaves. J Exp Bot 69: 891–903

15. Crawford NM, Campbell WH (1990) Fertile Fields. Plant Cell 2: 829–835

16. Czernic P, Gully D, Cartieaux F, Moulin L, Guefrachi I, Patrel D, Pierre O, Fardoux J, Chaintreuil C, Nguyen P, et al (2015) Convergent Evolution of Endosymbiont Differentiation in Dalbergioid and Inverted Repeat-Lacking Clade Legumes Mediated by Nodule-Specific Cysteine-Rich Peptides. Plant Physiol 169: 1254–1265

17. Dean JV, Harper JE (1988) The Conversion of Nitrite to Nitrogen Oxide(s) by the Constitutive NAD(P)H-Nitrate Reductase Enzyme from Soybean. Plant Physiol 88: 389–395

18. Douglas P, Morrice N, MacKintosh C (1995) Identification of a regulatory phosphorylation site in the hinge 1 region of nitrate reductase from spinach (Spinacea oleracea) leaves. FEBS Lett 377: 113–117

19. El-Gebali S, Mistry J, Bateman A, Eddy SR, Luciani A, Potter SC, Qureshi M, Richardson LJ, Salazar GA, Smart A, et al (2019) The Pfam protein families database in 2019. Nucleic Acids Res 47: D427–D432

20. Faure J-D, Vincentz M, Kronenberger J, Caboche M (1991) Co-regulated expression of nitrate and nitrite reductases. Plant J 1: 107–113

21. Feenstra WJ, Jacobsen E (1980) Isolation of a nitrate reductase deficient mutant of Pisum sativum by means of selection for chlorate resistance. Theor Appl Genet 58: 39–42

22. del Giudice J, Cam Y, Damiani I, Fung-Chat F, Meilhoc E, Bruand C, Brouquisse R, Puppo A, Boscari A (2011) Nitric oxide is required for an optimal establishment of the Medicago truncatula–Sinorhizobium meliloti symbiosis. New Phytol 191: 405–417

23. Guindon S, Dufayard J-F, Lefort V, Anisimova M, Hordijk W, Gascuel O (2010) New Algorithms and Methods to Estimate Maximum-Likelihood Phylogenies: Assessing the Performance of PhyML 3.0. Syst Biol 59: 307–321

24. Gupta KJ, Stoimenova M, Kaiser WM (2005) In higher plants, only root mitochondria, but not leaf mitochondria reduce nitrite to NO, in vitro and in situ. J Exp Bot 56: 2601–2609

25. Han R, Li C, Rasheed A, Pan X, Shi Q, Wu Z (2022) Reducing phosphorylation of nitrate reductase improves nitrate assimilation in rice. J Integr Agric 21: 15–25

26. Hardy RWF, Holsten RD, Jackson EK, Burns RC (1968) The Acetylene-Ethylene Assay for N2 Fixation: Laboratory and Field Evaluation 1. Plant Physiol 43: 1185–1207

27. Hoang DT, Chernomor O, von Haeseler A, Minh BQ, Vinh LS (2018) UFBoot2: Improving the Ultrafast Bootstrap Approximation. Mol Biol Evol 35: 518–522

28. Hoff T, Truong H-N, Caboche M (1994) The use of mutants and transgenic plants to study nitrate assimilation. Plant Cell Environ 17: 489–506

29. Horchani F, Prévot M, Boscari A, Evangelisti E, Meilhoc E, Bruand C, Raymond P, Boncompagni E, Aschi-Smiti S, Puppo A, et al (2011) Both plant and bacterial nitrate reductases contribute to nitric oxide production in medicago truncatula nitrogen-fixing nodules. Plant Physiol 155: 1023–1036

30. Kaiser WM, Huber SC (2001) Post-translational regulation of nitrate reductase: mechanism, physiological relevance and environmental triggers. J Exp Bot 52: 1981–1989

31. Kalyaanamoorthy S, Minh BQ, Wong TKF, von Haeseler A, Jermiin LS (2017) ModelFinder: fast model selection for accurate phylogenetic estimates. Nat Methods 14: 587–589

32. Kanamaru K, Wang R, Su W, Crawford NM (1999) Ser-534 in the hinge 1 region of Arabidopsis nitrate reductase is conditionally required for binding of 14-3-3 proteins and in vitro inhibition. J Biol Chem 274: 4160–4165

33. Kanayama Y, Kimura K, Nakamura Y, Ike T (1999) Purification and characterization of nitrate reductase from nodule cytosol of soybean plants. Physiol Plant 105: 396–401

34. Kato K, Okamura Y, Kanahama K, Kanayama Y (2003) Nitrate-independent expression of plant nitrate reductase in Lotus japonicus root nodules. J Exp Bot 54: 1685–1690

35. Katoh K, Standley DM (2013) MAFFT Multiple Sequence Alignment Software Version 7: Improvements in Performance and Usability. Mol Biol Evol 30: 772–780

36. Kolbert Z, Barroso JB, Brouquisse R, Corpas FJ, Gupta KJ, Lindermayr C, Loake GJ, Palma JM, Petřivalský M, Wendehenne D, et al (2019) A forty year journey: The generation and roles of NO in plants. Nitric Oxide - Biol Chem 93: 53–70

37. Kolbert Z, Ortega L, Erdei L (2010) Involvement of nitrate reductase (NR) in osmotic stress-induced NO generation of Arabidopsis thaliana L. roots. J Plant Physiol 167: 77–80

38. Lambeck IC, Fischer-Schrader K, Niks D, Roeper J, Chi J-C, Hille R, Schwarz G (2012) Molecular Mechanism of 14-3-3 Protein-mediated Inhibition of Plant Nitrate Reductase. J Biol Chem 287: 4562–4571

39. Lambert I, Pervent M, Le Queré A, Clément G, Tauzin M, Severac D, Benezech C, Tillard P, Martin-Magniette M-L, Colella S, et al (2020) Responses of mature symbiotic nodules to the whole-plant systemic nitrogen signaling. J Exp Bot 71: 5039–5052

40. Lea US, ten Hoopen F, Provan F, Kaiser WM, Meyer C, Lillo C (2004) Mutation of the regulatory phosphorylation site of tobacco nitrate reductase results in high nitrite excretion and NO emission from leaf and root tissue. Planta 219: 59–65

41. Lee T-Y, Bretaña NA, Lu C-T (2011) PlantPhos: using maximal dependence decomposition to identify plant phosphorylation sites with substrate site specificity. BMC Bioinformatics 12: 261

42. Letunic I, Bork P (2021) Interactive Tree Of Life (iTOL) v5: an online tool for phylogenetic tree display and annotation. Nucleic Acids Res 49: W293–W296

43. Li W, Godzik A (2006) Cd-hit: a fast program for clustering and comparing large sets of protein or nucleotide sequences. Bioinforma Oxf Engl 22: 1658–1659

44. Lillo C, Kazazaic S, Ruoff P, Meyer C (1997) Characterization of Nitrate Reductase from Light-and Dark-Exposed Leaves (Comparison of Different Species and Effects of 14-3-3 Inhibitor Proteins). Plant Physiol 114: 1377–1383

45. Lillo C, Lea US, Leydecker M-T, Meyer C (2003) Mutation of the regulatory phosphorylation site of tobacco nitrate reductase results in constitutive activation of the enzyme in vivo and nitrite accumulation. Plant J Cell Mol Biol 35: 566–573

46. Lillo C, Meyer C, Lea US, Provan F, Oltedal S (2004) Mechanism and importance of Post-translational regulation of nitrate reductase. J Exp Bot 55: 1275–1282

47. Lopez G, Ahmadi SH, Amelung W, Athmann M, Ewert F, Gaiser T, Gocke MI, Kautz T, Postma J, Rachmilevitch S, et al (2023) Nutrient deficiency effects on root architecture and root-to-shoot ratio in arable crops. Front. Plant Sci. 13:

48. Meyer C, Stitt M (2001) Nitrate Reduction and signalling. *In* PJ Lea, J-F Morot-Gaudry, eds, Plant Nitrogen. Springer, Berlin, Heidelberg, pp 37–59

49. Minh BQ, Schmidt HA, Chernomor O, Schrempf D, Woodhams MD, von Haeseler A, Lanfear R (2020) IQ-TREE 2: New Models and Efficient Methods for Phylogenetic Inference in the Genomic Era. Mol Biol Evol 37: 1530–1534

50. Opperdoes FR, Lemey P (2009) Phylogenetic analysis using protein sequences. *In* A-M Vandamme, M Salemi, P Lemey, eds, Phylogenetic Handb. Pract. Approach Phylogenetic Anal. Hypothesis Test., 2nd ed. Cambridge University Press, Cambridge, pp 313–342

51. Pelsy F, Caboche M (1992) Molecular Genetics of Nitrate Reductase in Higher Plants. *In* JG Scandalios, TRF Wright, eds, Adv. Genet. Academic Press, pp 1–40

52. Planchet E, Gupta KJ, Sonoda M, Kaiser WM (2005) Nitric oxide emission from tobacco leaves and cell suspensions: Rate limiting factors and evidence for the involvement of mitochondrial electron transport. Plant J 41: 732–743

53. Postgate JR (1982) Biology Nitrogen Fixation: Fundamentals. Philos Trans R Soc Lond B Biol Sci 296: 375–385

54. Puppo A, Pauly N, Boscari A, Mandon K, Brouquisse R (2013) Hydrogen peroxide and nitric oxide: key regulators of the Legume-Rhizobium and mycorrhizal symbioses. Antioxid Redox Signal 18: 2202–2219

55. Rancurel C, van Tran T, Elie C, Hilliou F (2019) SATQPCR: Website for statistical analysis of real-time quantitative PCR data. Mol Cell Probes 46: 101418

56. Ratet P, Wen J, Cosson V, Tadege M, Mysore KS (2010) Tnt1 Induced Mutations in Medicago: Characterization and Applications. Handb. Plant Mutat. Screen. John Wiley & Sons, Ltd, pp 83–99

57. Rockel P, Strube F, Rockel A, Wildt J, Kaiser WM (2002) Regulation of nitric oxide (NO) production by plant nitrate reductase in vivo and in vitro. J Exp Bot 53: 103–110

58. Ryan SA, Nelson RS, Harper JE (1983) Soybean Mutants Lacking Constitutive Nitrate Reductase Activity. Plant Physiol 72: 510–514

59. Sakihama Y, Nakamura S, Yamasaki H (2002) Nitric oxide production mediated by nitrate reductase in the green alga Chlamydomonas reinhardtii: an alternative NO production pathway in photosynthetic organisms. Plant Cell Physiol 43: 290–297

60. Salas A, Cabrera JJ, Jiménez-Leiva A, Mesa S, Bedmar EJ, Richardson DJ, Gates AJ, Delgado MJ (2021) Bacterial nitric oxide metabolism: Recent insights in rhizobia. Adv Microb Physiol 78: 259–315

61. Sánchez C, Gates AJ, Meakin GE, Uchiumi T, Girard L, Richardson DJ, Bedmar EJ, Delgado MJ (2010) Production of nitric oxide and nitrosylleghemoglobin complexes in soybean nodules in response to flooding. Mol Plant Microbe Interact 23: 702–711

62. Silveira JAG, Matos JCS, Cecatto VM, Viegas RA, Oliveira JTA (2001) Nitrate reductase activity, distribution, and response to nitrate in two contrasting *Phaseolus* species inoculated with *Rhizobium* spp. Environ Exp Bot 46: 37–46

63. Slater GSC, Birney E (2005) Automated generation of heuristics for biological sequence comparison. BMC Bioinformatics 6: 31

64. Streeter JG (1985a) Nitrate Inhibition of Legume Nodule Growth and Activity 1: I. Long Term Studies with a Continuous Supply of Nitrate. Plant Physiol 77: 321–324

65. Streeter JG (1985b) Nitrate Inhibition of Legume Nodule Growth and Activity 1: II. Short Term Studies with High Nitrate Supply. Plant Physiol 77: 325–328

66. Su W, Huber SC, Crawford NM (1996) Identification in vitro of a post-translational regulatory site in the hinge 1 region of Arabidopsis nitrate reductase. Plant Cell 8: 519–527

67. Tadege M, Wen J, He J, Tu H, Kwak Y, Eschstruth A, Cayrel A, Endre G, Zhao PX, Chabaud M, et al (2008) Large-scale insertional mutagenesis using the Tnt1 retrotransposon in the model legume Medicago truncatula. Plant J Cell Mol Biol 54: 335–347

68. Terpolilli JJ, Hood GA, Poole PS (2012) What Determines the Efficiency of N2-Fixing Rhizobium-Legume Symbioses? Adv Microb Physiol. doi: 10.1016/B978-0-12-398264-3.00005-X

69. Udvardi MK, Day DA (1997) METABOLITE TRANSPORT ACROSS SYMBIOTIC MEMBRANES OF LEGUME NODULES. Annu Rev Plant Biol 48: 493–523

70. Wang D, Zeng S, Xu C, Qiu W, Liang Y, Joshi T, Xu D (2017) MusiteDeep: a deep-learning framework for general and kinase-specific phosphorylation site prediction. Bioinformatics 33: 3909–3916

71. Wang P, Du Y, Li Y, Ren D, Song C-P (2010) Hydrogen Peroxide–Mediated Activation of MAP Kinase 6 Modulates Nitric Oxide Biosynthesis and Signal Transduction in Arabidopsis[w]. Plant Cell 22: 2981–2998

72. Wang R, Tischner R, Gutiérrez RA, Hoffman M, Xing X, Chen M, Coruzzi G, Crawford NM (2004) Genomic analysis of the nitrate response using a nitrate reductase-null mutant of arabidopsis. Plant Physiol 136: 2512–2522

73. Wilkinson JQ, Crawford NM (1991) Identification of the Arabidopsis CHL3 gene as the nitrate reductase structural gene NIA2. Plant Cell 3: 461–471

74. Wilkinson JQ, Crawford NM (1993) Identification and characterization of a chlorate-resistant mutant of Arabidopsis thaliana with mutations in both nitrate reductase structural genes NIA1 and NIA2. MGG Mol Gen Genet 239: 289–297

75. Wojciechowski MF, Lavin M, Sanderson MJ (2004) A phylogeny of legumes (Leguminosae) based on analysis of the plastid matK gene resolves many well-supported subclades within the family. Am J Bot 91: 1846–1862

76. Yamasaki H, Sakihama Y (2000) Simultaneous production of nitric oxide and peroxynitrite by plant nitrate reductase: in vitro evidence for the NR-dependent formation of active nitrogen species. FEBS Lett 468: 89–92

